# Transposable Elements and piRNAs interaction prediction with Predictive Bi-Clustering Trees

**DOI:** 10.1101/2024.02.28.582449

**Authors:** Hiago Freire Oliveira, Renato A. C. dos Santos, Ricardo Cerri

## Abstract

PIWI-interacting RNAs (piRNAs) are a class of noncoding RNAs whose actions range from regulating gene expression to silencing Transposable Elements, characterized for being from 21 to 35 nucleotides long, displaying a uracil bias at the 5’ end, and a 2’-O-methylation at the 3’ end. Transposable Elements (TEs) are genetic elements that move within host genomes. TE replication can promote harmful recombination events by generating breaks in DNA double strands, in addition to interfering with expression. Silencing of these elements by piRNAs occurs in the germ line in most animals and is essential for maintaining genome integrity. In this work, the problem of in silico interaction prediction between piRNAs and TEs was addressed by a decision tree-based algorithm, namely Predictive Bi-Clustering Trees (PBCT). In order to improve the algorithm’s performance, the piRNA-TE interaction matrix was reconstructed using a Beta-distribution-rescored Neighborhood Regularized Logistic Matrix Factorization (NRLMFβ) algorithm. PBCT was tested in 5-fold and 10-fold cross-validation configurations, both with the original interaction matrix (BICT) and the interaction matrix reconstructed by NRLMFβ (BICTR). Although not being able to predict positive interactions satisfactorily given the huge dataset imbalance, advantages could be observed when using matrix factorization. Comparatively, in the BICT method, PBCT presented higher values of AUROC and AUPRC. However, in the BICTR method, PBCT was able to correctly predict more positive interactions, which are, in fact, the primary interest of this study. Potential biological applications and ways to improve the algorithm’s performance were also discussed.

**Author summary:** piRNAs and transposable elements are biomolecules that interact in the germ lime in most animals, such that piRNAs silence these elements to keep genome integrity. However, detecting which piRNA interacts with which TE is a laborious task with low results, given that the rules that govern these interactions still need to be fully elicited. In this paper, we addressed the interaction prediction pair piRNA-TE using a multi-label decision-tree-like algorithm called PBCT applied to *in vivo* known interactions. Given that it is a Positive-Unlabeled Learning problem, since we cannot be sure of a biological negative interaction, we reconstructed the interaction matrix employing an NRLMFβ algorithm. We compared the results given the original interaction matrix and the reconstructed matrix. The results with this algorithm and parameters could have been better, even though the reconstruction has proven fruitful. Further, we addressed our problem with other multi-label learning approaches and briefly compared them. We also discussed potential biological applications and ways to improve the algorithm’s performance.

## Introduction

### PIWI-interacting RNAs

RNA interference was first described in *Caenorhabditis elegans*, a model nematode, as a means to manipulate gene expression [1]. This mechanism works by noncoding RNA molecules that can regulate genome function, generally with inhibitory effects, in different levels, such as chromosome segregation, chromatin structure, translation, and transcription [2]. A class of regulatory RNA that integrates RNA interference is the miRNA (microRNA) class. The first discovered miRNAs were *lin-4* [3] and *let-7* [4], both acting in *C. elegans* heterochronic gene pathway, that is, the timing regulation of events in development. These molecules have a regulatory function and are approximately 22 nucleotides long. miRNA sequences are arranged throughout the human genome, whether in exonic, intronic, or even intergenic regions [5]. Moreover, misexpression of miRNAs usually leads to phenotypes that resemble loss-of-function for their targets, which makes their regulation very tight by their very own targets [2].

Another class of regulatory RNA is the siRNA (small interfering RNA) class. Several aspects make them similar to miRNAs, such as the approximate 22 nucleotide length, biogenesis from dsRNA (double-strand DNA), processing by the cytoplasmic enzyme Dicer, and loading by AGO enzymes. The siRNAs are loaded by the AGO component of the RNA-induced silencing complex (RISC), which is guided to the complementary mRNA molecule, resulting in its cleavage and degradation [6]. However, there are distinctions since miRNAs act, in general, by partial base pairing with the target mRNA and restrict the expression of several genes of similar sequences, while siRNAs act by perfect pairing and are specific to regions of the target mRNA. Not only, while miRNAs have a role more related to the regulation of gene expression, siRNAs appear to be related to the recognition of nucleic acids for immunological purposes and can be used, by biotechnological applications, against all types of viral genomes, whether DNA or RNA, whether double or single stranded [7].

PIWI-interacting RNAs (piRNAs) comprise a class of regulatory RNAs defined by the specificity of loading by PIWI (P-element-induced wimpy testis) proteins, present predominantly in the cell nucleus [5], in addition to being processed from long single-stranded precursor transcripts and not requiring processing by Dicer. piRNAs range from 21 to 35 nucleotides, have a uracil bias at the 5’ end, and carry a 2’-O-methylation at the 3’ end. Genomic loci called piRNA clusters are piRNA precursors. The effects of piRNA action range from regulating gene expression to fighting viral infections and silencing Transposable Elements. Similar to RISC, piRNAs guide PIWI proteins to target RNA for cleavage. In addition, they promote the assembly of heterochromatin and DNA methylation [8].

Ago, PIWI, WAGO (worm-specific AGO), and Trypanosoma Ago families comprise the major classifications of eukaryotic Argonaut proteins. The architecture of the Argonaut proteins comprises a basic nucleus constituted by the N-terminal (amino-terminal), PAZ (PIWI-Argonaute-Zwille), MID (Middle), and PIWI domains. Structural differences between Argonaut proteins appear to be closely related to the types of small RNA with which they bind and the strategies adopted to bind to their targets [9].

In *C. elegans*, piRNAs are classically known as 21U-RNAs, given the uracil bias and the persistent length of 21 nucleotides. Unlike the piRNA processing mechanism from transcripts of a long single-stranded precursor discussed earlier, type I 21U-RNAs are transcribed from approximately 12,000 minigenes by the transcription factor of the Forkhead family of proteins, which binds to the Ruby motif upstream to each piRNA precursor. A 5’-capped precursor transcript of only 25-27 nucleotides is generated from each type I minigene. The precursor is loaded into PRG-1 protein (P53-Responsive Gene 1) and processed into mature piRNA. Type II 21U-RNAs are transcribed from full-length protein-producing gene promoters [8].

One recognized important role of piRNAs in the nematode is the ability to recognize self and non-self transcripts. This allows the recognition and silencing of transgenes and new transposonic insertions. Unlike cytoplasmic PIWI proteins from other animals, the slicing ability of the *C. elegans* PIWI protein (PRG-1) is not imperative, as piRNAs induce transcription of secondary siRNAs by an RNA-dependent RNA polymerase using the target itself as a template. The resulting siRNAs are called 22G-RNA (due to first nucleotide bias and transcript length). 22G-RNAs are loaded into WAGO proteins, which ultimately silence the non-self transcript [8]. It is equally notable that certain AGOs, such as WAGO-9 and CSR-1 (Chromosome Segregation and RNAi Deficient 1), are involved in epigenetic inheritance processes such as silencing non-self transcripts and protecting self transcripts, respectively [10].

Most of the approximately 15,000 piRNAs encoded by the nematode do not show broad complementarity to Transposons. Many endogenous genes contain target sites for piRNAs but exhibit resistance to silencing, mainly due to the presence of PATCs (Periodic A_n_/T_n_ Clusters) [11]. PATCs are non-coding elements spaced by ten base pairs, so they can enable germline carrier gene expression in repressive chromatin domains, thus representing the basis of the cellular immune system [12]. The seed region of a piRNA spans from the second to the seventh nucleotide and is distinguished by optimal pairing to target sites. Outside this region, up to six unpaired bases make the transcript a potential piRNA target [11].

### Transposable elements and piRNAs

Transposable Elements (TEs) are genetic elements capable of moving within a host genome [13]. About 12% of the genome of *C. elegans* is composed of these elements [14]. TEs are classified into two major classes: class 1, made up of Retrotransposons, and class 2, made up of DNA Transposons. Retrotransposons carry out the transposition from a genomic copy that is transcribed to an RNA intermediate, which is reverse transcribed to DNA by a reverse transcriptase (“copy-and-paste” mechanism) [15]. Notable examples of this class are the LTR (Long Terminal Repeats), among which is the gypsy family, which is controlled by the flamenco cluster in *Drosophila* [16], and the Cer group (composed of gypsy and Bel elements), which makes up about 0.4% of the entire genome of *Caenorhabditis elegans* [17].

DNA Transposons are divided into two subclasses: subclass 1, comprising TEs that have terminal inverted repeats (TIR) and that use the “cut-and-paste” mechanism, and subclass 2, comprising TEs that replicate without double-stranded DNA cleavage (“copy-and-paste”), differing from Retrotransposons by the absence of RNA intermediates, in such a way that a single strand of DNA is displaced [15]. Notable examples of subclass 1 are the P elements, associated with the cause of hybrid dysgenesis in *Drosophila* [18] and the Tc1/mariner, first discovered in *C. elegans*, but which are probably the most pervasive DNA Transposons in nature [19]. Subclass 2 examples worthy of notice, a much less diverse subclass, are the Helitrons, which are transposed via replication by a rolling circle and which, although better characterized in plants, constitute about 2% of the genome of the model nematode [20].

Replication of TEs can promote harmful recombinational events by generating breaks in the DNA double-strand, in addition to interference in expression, since their promoters can lead to aberrant transcription of neighboring genes. However, Transposons must integrate into the germ cell’s DNA to survive. In most animals, at least a subset of piRNAs defend the germline genome against TE mobilization. Once silenced, mutations can ultimately inactivate proteins encoded by Transposons, leading to their downfall [8].

The presence of piRNAs is not restricted to the germline in all animals. Most arthropods possess, in addition, piRNAs in somatic cells, presenting mechanisms more diverse than the strict silencing of TEs, such as defense against viruses (in addition to siRNAs) in *Aedes aegypti*. The role of defense against Transposons in ancestral arthropods was probably dependent not on piRNAs but on siRNAs catalyzed by an RNA-dependent RNA polymerase (as observed in *C. elegans*), which provided dsRNA precursors different from those of RNA polymerase II, that is, a new range of substrate diversity. In this sense, the redefinition of the role of piRNAs in the evolutionary history of arthropods (such as *Drosophila*) is evidenced by the diversity of functions of somatic piRNAs, which are considered to be ancestral to all arthropods [21].

### Interaction Prediction

The cross-linking, ligation, and sequencing of hybrids (CLASH) protocol provides methods for discovering RNA-RNA interactions *in vivo* so that work is eased and prior knowledge of the molecular pairs is not necessary, differing from methods such as X-ray crystallography and nuclear magnetic resonance which require knowing the pairs.

CLASH uses UV light cross-linking methods, followed by affinity purification of RNA-protein complexes and binding and sequencing of RNA-RNA hybrids. These sequenced chimeric molecules are the final product of CLASH, so they represent a binding event between the small RNA and a fragment of its target mRNA [22].

Interactions of piRNA-mRNA kind were verified *in vivo* in *C. elegans* employing CLASH protocol, so that it was observed that all germline mRNAs are subject to piRNA surveillance, that is, the entire transcriptome. Besides, interactions between piRNAs and other ncRNAs (non-coding RNAs), although much less frequent, were verified, of special relevance piRNA-tRNA interactions (transfer RNA), since in *Drosophila* the accumulation of unprocessed tRNA due to a mutation leads to failure of PIWI-mediated Transposon silencing [23].

Considering the already discussed properties of piRNA targets (such as the limit of unpaired bases and seed sequence) and the high cost of experiments in physical laboratories, it is unsurprising that computational tools have emerged to provide data concerning piRNA targets in different species. pirScan, for example, is a web-based tool suitable for identifying target sites of *Caenorhabditis elegans* piRNAs, given a mRNA or an intron-free DNA sequence [24]. piRTarBase is an interactive online database that provides piRNA interaction sites in *C. elegans* and *C. briggsae*, both predicted and experimentally verified, from gene input. Also, mRNA targets can be obtained from a piRNA input [25].

However, an *in silico* prediction model of interactions between piRNAs and TEs exclusively was not found. Extracting useful knowledge from these data falls into a major problem of computational biology, which requires the development of case-specific tools [26]. Since TEs are the elements that, in fact, are affected by piRNAs, the interest in understanding how a piRNA recognizes a Transposable Element is definitive, even given the knowledge of the existence of PATCs. Thus, computer modeling can help to find patterns that are difficult to observe in direct experimentation.

Due to the complex nature of this mechanism, sequence alignment may not be sufficient to predict target interactions. In this way, machine learning has proved to be an excellent tool for problem resolution in several biological circumstances [26]. Machine learning can be defined as the area of knowledge interested in studying algorithms that allow computer programs to automatically improve through experience, that is, without explicitly being programmed. In prediction problems, different performance criteria can be optimized, such as precision and accuracy [27].

The problem of predicting the interaction between two pairs of objects can be approached through decision trees [28]. Decision trees are one of the simplest yet most effective machine learning techniques. They aim to classify unknown objects being built by analyzing a set of known objects. Classification is based on questions about attributes associated with items. A tree node, that being the root or internal nodes, poses attribute test conditions to split the observation attribute’s value. The terminal node, the one that has no outgoing edges, is called the leaf node. The object is, therefore, classified according to the class label associated with the leaf node [29]. A tree node corresponds to a question, which can follow to lower nodes (two or more) depending on the answer, that is, the value of the observation attribute. The final node, the one that asks no more questions, is called the leaf, which is the class of the object [30].

A powerful approach to building decision trees is Predictive Clustering Trees (PCTs) [31], so each node is considered a cluster of data, building up the tree inductively from top to bottom. Thus, the total dataset is the root node cluster, from which new nodes are recursively split by testing one of the features. The best division is obtained by considering all the characteristics’ division points and evaluating them by the information gain criterion. A stop criterion is based on the purity of the node so that when the data contained in that same node are pure in relation to the target, the prediction is given according to the majority class in the node. The entropy function is usually used as a measure of the purity of the nodes so that the information gain is the difference between the entropy of the distribution of the classes in the parent node and the weighted average of the entropy of the resulting children nodes [30]. Since a piRNA can interact with more than one Transposable Element and vice versa, multi-label classification can approach the problem, considering that objects can be associated with several classes simultaneously. In this way, PCTs can predict multiple targets simultaneously, thus characterizing a decision tree with multiple outputs [28].

Another way of predicting interactions is to work with a matrix of interactions, where each row is a vector of features that numerically depict the piRNA, and each column is a vector of features that numerically depict the TE. In this sense, *e*.*g*. if piRNA 3 is known to interact solely with TE 7, the third row (feature vector of piRNA 3) will record **1** in the seventh column (feature vector of TE 7), while **0** in the remaining columns. In this case, algorithms that can work with both feature spaces are desirable. In this direction, PBCT (Predictive Bi-Clustering Trees) is an algorithm that, as a bi-clustering algorithm, makes the simultaneous grouping of rows and columns [32]. Therefore, unlike the PCT algorithm, which works with only one set of features, PBCT builds a decision tree by partitioning both feature spaces (row and column objects) [33]. Therefore, considering that both piRNA and TE information are required to predict the interaction, PBCT is suitable for the present study.

The remainder of this paper is organized as follows. We first introduce how the interaction matrix was built, how it was reconstructed to alleviate the imbalance of classes, and how the prediction method works and is evaluated. Then, we present and discuss the results of both original and reconstructed matrix prediction methods, pointing to the real implications of their usage. Lastly, we present and discuss the results of alternative approaches to this problem.

## Materials and methods

### Biological interactions

The piRNA-TE interactions used in the present study were previously described by Shen *et al*. [23], who highlighted detailed instructions regarding the discovery of *in vivo* piRNA-target interactions through CLASH in *Caenorhabditis elegans*. Of all interactions, 19,092 reads in totality (with redundancy) point to the piRNA-TE interaction. These readings also indicate the precise sites of interaction of the molecules.

The additional information from the piRNA-TE reads of CLASH was removed, so only the bases were retained. A table was generated from these interactions so that the piRNA sequences composed the row index and the TE sequences composed the column index. The table was populated with 0 and 1 so that each piRNA-TE interaction was represented by 1 and unknown pairs (might or not interact) by 0. Thus, a binary interaction matrix was generated. Once built, the indices and matrix were exported independently in 3 files: the binary matrix as a CSV (comma-separated values) file and the indices as two individual FASTA files. A total of 5,218 (unique) interactions were counted in a matrix of 13,841,400 items (approximately 0.000377% of the total), characterizing a very sparse dataset.

### Feature generation

FASTA files were used to extract features from sequences using the *Pse-in-One* software [34]. Since the prediction task cannot deal with strings, the sequences must be converted into fixed-dimensional vectors (holding 18 features, in the case of this study) containing the key features. Each vector is called a feature vector. *Pse-in-One* has three web sub-servers, among which *PseDAC-General* (pseudo deoxyribonucleic acid compositions for DNA sequences) was used. *PseDAC-General* contains three categories, namely nucleotide composition, nucleotide autocorrelation, and pseudonucleotide composition, such that 16 modes are distributed among these categories [34]. The features calculated in the present study derive from the *PseDNC* (Pseudo dinucleotide composition) mode, which is part of the pseudonucleotide composition category. The PseDNC mode considers local structural parameters of the sequence, that is, angular and translational parameters [35]. The fixed-dimension vectors, given by PseDNC, are computed by:

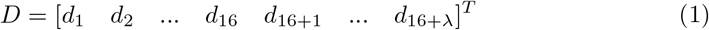

so that:

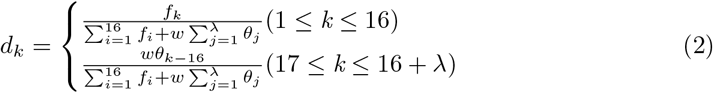

given *f*_*k*_ (k = 1,2,3,…,16), that is the normalized occurrence frequency of the *k*^*th*^ dinucleotide, and *θ*_*j*_ (j = 1,2,3,…,*λ*), that is the *j*^*th*^-tier correlation factor, reflecting the sequence-order correlation between the *j*^*th*^ most contiguous dinucleotide. The parameters *λ* (an integer) and *w* (ranging from 0 to 1) stand for, respectively, the highest counted rank of the correlation in the sequence and the weight factor. Higher *λ* values lead to incorporating more sequence-order effects. Therefore, the DNA sequence can be denoted as a (16+*λ*) dimensional vector rather than a 16-dimensional one, as would be expected if it was a contiguity frequency vector (4^2^ nucleic acids). For a full exposition of the method, consult Chen *et al*. [35].

The software then generated two new .csv files, one of which is the file with the feature vectors extracted from each of the piRNAs, and the other is the file with features extracted from each of the TEs in the exact order in the previously generated binary matrix. Figure 1 represents the generated files and their organization for the interaction prediction task. We can see that a piRNA can interact with two or more TEs simultaneously, characterizing a multi-label task. Three .csv files were generated: i) one with piRNAs features, ii) one with TEs features, iii) and another with information on interactions between pairs of piRNAs and TEs. These files were used in the PBCT algorithm for multi-label interaction prediction [28].

**Fig 1.**
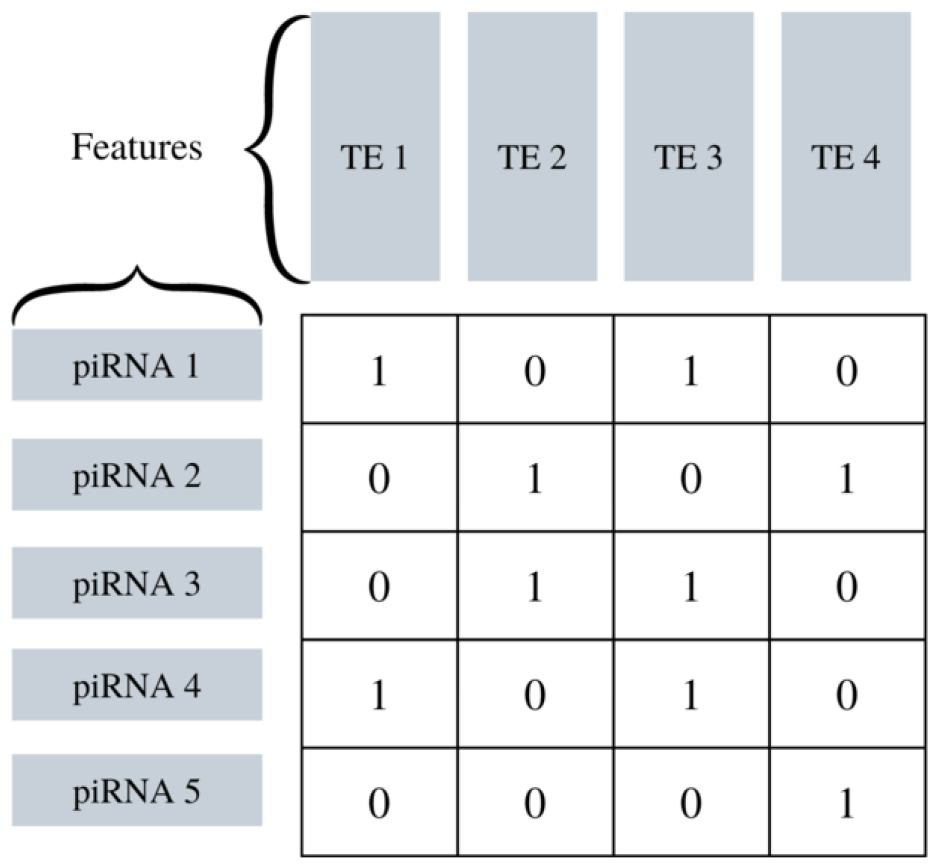
Example of a binary matrix of interactions between piRNAs and TEs, represented by their respective features. Adapted from [28].

According to Pliakos & Vens [36], building learning models through ensemble-type trees (that is, the combination of several decision trees in a single classifier), when elaborated from the reconstruction of the learning space output, leads to better prediction results. Thus, the authors proposed the Bi-Clustering Trees with output space Reconstruction (BICTR). The reconstruction of the output space is performed with NRLMF (Neighborhood Regularized Logistic Matrix Factorization).

### NRLMF

Let the pairs R and E be two finite sets of nodes *R* = {*r*_1_, …, *r*_|*R*|_}, corresponding to piRNAs, and *E* = {*e*_1_, …, *e*_|*E*|_}, corresponding to TEs [36]. Given a binary interaction matrix *Y* ∈ ℝ ^|*R*|*×*|*E*|^, one can compute the factorization by calculating matrices *U* ∈ ℝ ^|*R*|*×k*^ and *V* ∈ ℝ ^|*E*|*×k*^, such that the matrices *U* and *V* are considered k-dimensional latent representations of piRNAs and TEs, respectively. The *UV* ^*T*^ product should be approximately *Y* [37].

The nodes in the pair of sets R and E are sequence similarity vectors so that a similarity vector of a molecule is defined by a real vector with values between 0 and 1. The set with similarity vectors is used solely to reconstruct the output space, given that the original output space is the interaction matrix of the piRNAs features and the TEs features, calculated by *Pse-in-One*, as mentioned above. We computed the similarities between the sequences of both piRNAs and TEs with normalized Smith-Waterman score, which is given by:

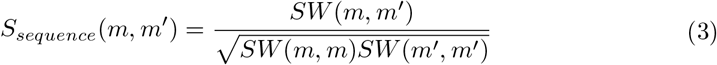

for sequences m and m’, where *SW* (*m, m*^*′*^) stands for the Smith-Waterman score [38]. The result is two sets of sequence similarity vectors, one for the piRNAs sequences and another for the TEs sequences, as already mentioned. In order to illustrate the process, Figure 2 depicts a simple sequence similarity matrix for the Transposable Elements. As one can expect, the first row is identical to the first column, and so on.

**Fig 2.**
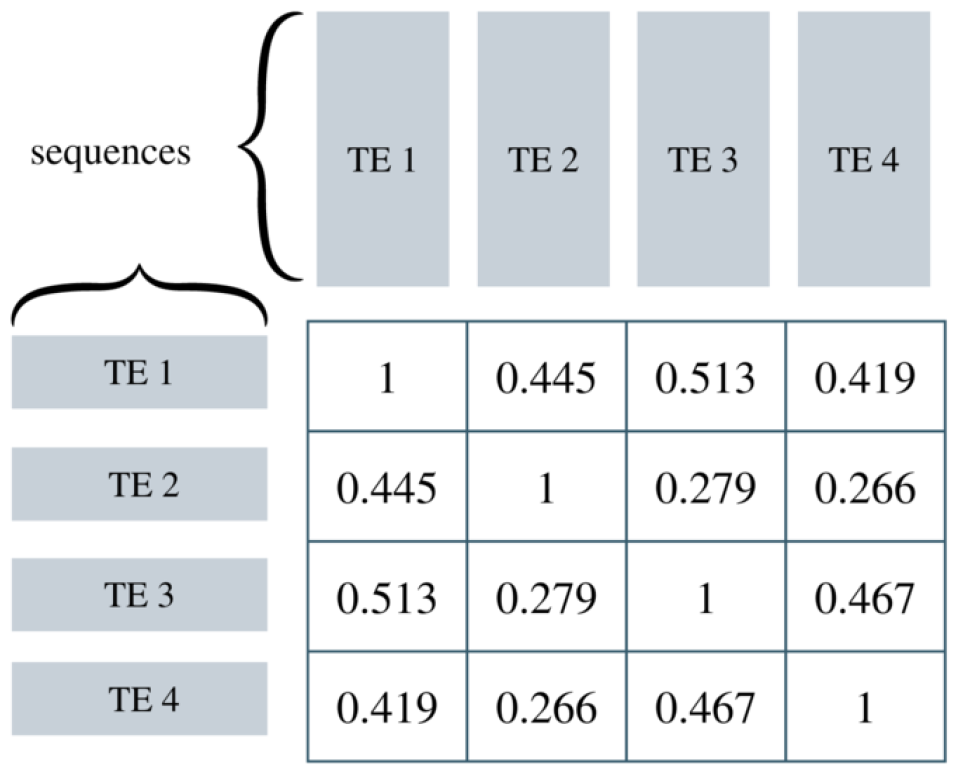
Example of a TE sequence similarity matrix.

In order to model the probability that piRNA *r*_*i*_ interacts with TE *e*_*j*_, *p*_*ij*_ is expressed as:

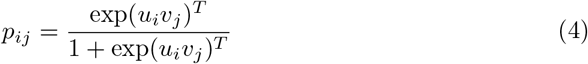

where *u*_*i*_ and *v*_*j*_ are k-dimensional vectors and latent representations of, respectively, *r*_*i*_ and *e*_*j*_. After the prediction and regularization model, an objective function is obtained, taking into account the hyperparameters of regularization and weighting of the observations in the optimization process. For a full understanding of the method and the complete derivation of the equations, the reader is referred to [34]. The reconstruction of the output space is relevant, given that in our piRNA-TE interaction prediction task, there are no truly negative interaction pairs (represented by 0). Instead, there are known positive interactions (represented by 1) and unlabeled interactions (since they simply may not have been observed yet in the laboratory). In this sense, this configuration is called Positive-Unlabeled Learning (or positive - not labeled). The reconstruction, therefore, aims to alleviate the imbalance of classes (considering that there are many more unlabeled ones than positive ones), taking into account that 0 values do not, in fact, typify negative interactions [36].

### NRLMF*β*

As noted by Ban *et al*.. [38], the prediction with NRLMF is not accurate when there is little information about the interaction pairs. Thus, they proposed the NRLMFβ (Beta-distribution-scored Neighborhood Regularized Logistic Matrix Factorization). This new score takes advantage of the fact that NRLMF is established on a statistical model based on the Bernoulli distribution and that the beta distribution is the conjugate prior to the Bernoulli distribution so that the latter can reflect the amount of information from interactions for its shape based on Bayesian inference [38]. The *s*_*ij*_ score of the NRLMF is calculated according to Equation 5.

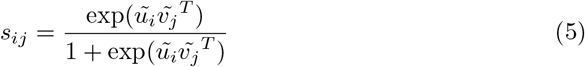

The *s*_*ij*_ score is used to generate a new matrix. This score takes the modified latent feature vectors 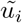 and 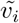, given by a recalculation of the vectors in the scenario of a molecule having no targets. *s*_*ij*_ becomes very low when there is little information about the interaction. The beta distribution, described by:

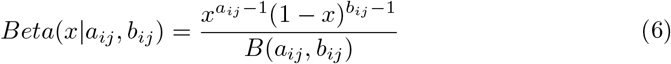

fixes this problem by means of the relational expression:

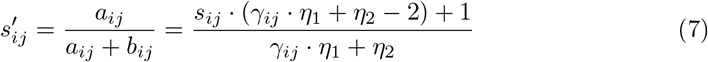

so that the parameters *a*_*ij*_ and *b*_*ij*_ are computed by 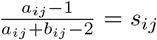 and *a*_*ij*_ + *b*_*ij*_ = *γ*_*ij*_ *· η*_1_ + *η*_2_. Thus, the reconstruction of the output space in our study was performed with NRLMFβto improve the prediction accuracy. We manually defined the hyperparameters *η*_1_ and *η*_2_, expressed in Equation 7, according to the authors’ suggestion. These hyperparameters are constants of the concentration equation of the beta distribution that concern the shape of the distribution. The complete derivation can be verified in [34] and [38]

A new matrix of interactions is generated with the reconstruction, denoted *Ŷ*. Each new item of the new matrix, such that *ŷ*_*ij*_ ∈*Ŷ*, is given by:

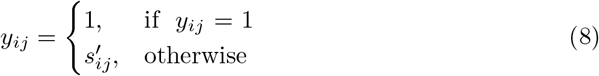

With the recalculated matrix, PBCT algorithm was applied, using the feature space generated from the piRNAs using Pse-in-One (*X1*), the feature space generated from the TEs using Pse-in-One (*X2*), and the new matrix of interactions (*Ŷ*) calculated by NRLMFβ. The PBCT applied to this new matrix is the previously named BICTR method (Figure 3). The PBCT algorithm was also applied using the original interaction matrix *Y* to compare performances. In contrast to BICTR, the PBCT applied to the original matrix was called BICT (Bi-Clustering Trees).

**Fig 3.**
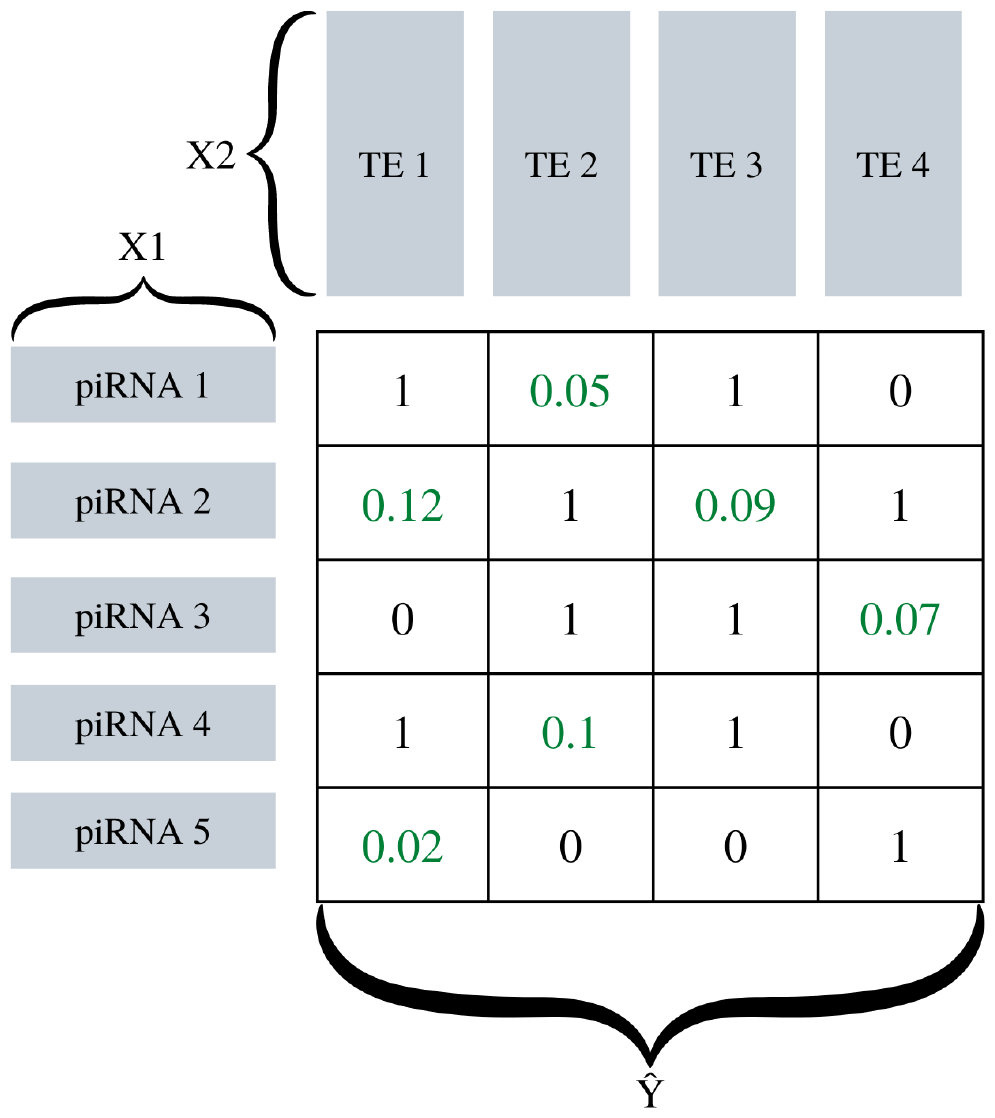
Example of input data for the PBCT algorithm in the BICTR methodology. The data without repopulation, that is, the original data, are the input data for the BICT methodology.

### Prediction

As presented here, the interaction prediction is a molecule-molecule interaction network inference. The prediction is a node pair classification type task, meaning it is a supervised learning application. The goal is to obtain the probability of interaction between two pairs of nodes. A learning model is built on top of TE-piRNA training set pairs. Once the learning process is completed, the model can predict unknown interaction pairs [36].

There are different approaches to predicting an interaction network, namely the local and global approaches. The local approach is based on dividing the data into *N* subsets, in accordance with the number of classes *N*, so that each subset has objects associated with one of the *N* classes, leading to the training of *N* binary classifiers. Integrating the *N* outputs leads to the prediction in the test objects. The global approach, on the other hand, is based on dealing with a learning method adjusted in order to be able to deal with all classes simultaneously, without dividing the problem into sub-tasks, using a single classifier [33]. If we collapse the feature spaces into rows, we can only yield a single output. However, as already described, PBCT uses rows and columns to predict a matrix, yielding a multiple output space.

As characterized before, considering *X1* the feature space generated from the piRNAs using Pse-in-One, *X2* the feature space generated from the TEs using Pse-in-One, and *Y* the interaction matrix, the local and global approaches can be illustrated by Figure 4.

**Fig 4.**
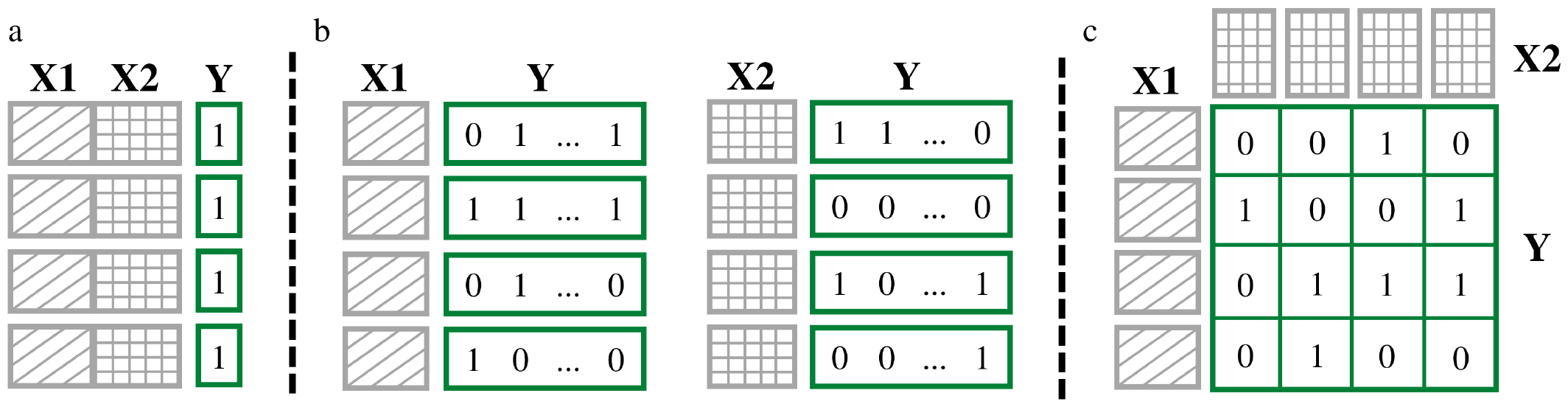
Interaction prediction approaches. In *a*, the global single-output method. In *b*, the local multiple-output method. In *c*, the global multiple-output method. It is the input data for PBCT, as already shown. Adapted from [28]

When evaluating an interaction prediction algorithm, four prediction tasks can be considered:

- Learning piRNAs (rows) - Learning TEs (columns) (*Lr × Lc*): a trivial task, as it consists in predicting interactions between piRNAs and TEs included in the learning procedure;
- Test piRNAs - Learning TEs (*Tr × Lc*): consists in predicting interactions between TEs included in the learning procedure and unknown piRNAs;
- Learning piRNAs - Test TEs (*Lr × Tc*): consists in predicting interactions between piRNAs included in the learning procedure and unknown TEs;
- Test piRNAs - Test TEs (*Tr × Tc*): consists in predicting interactions between unknown piRNAs and unknown TEs. Such task is, naturally, the more challenging one.

The prediction scheme of a piRNA-TE interaction network, comprising the above tasks, is illustrated in Figure 5. As one can see, Lr × Lc task encompass the largest portion of the interaction matrix, while Tr × Tc the smallest one.

**Fig 5.**
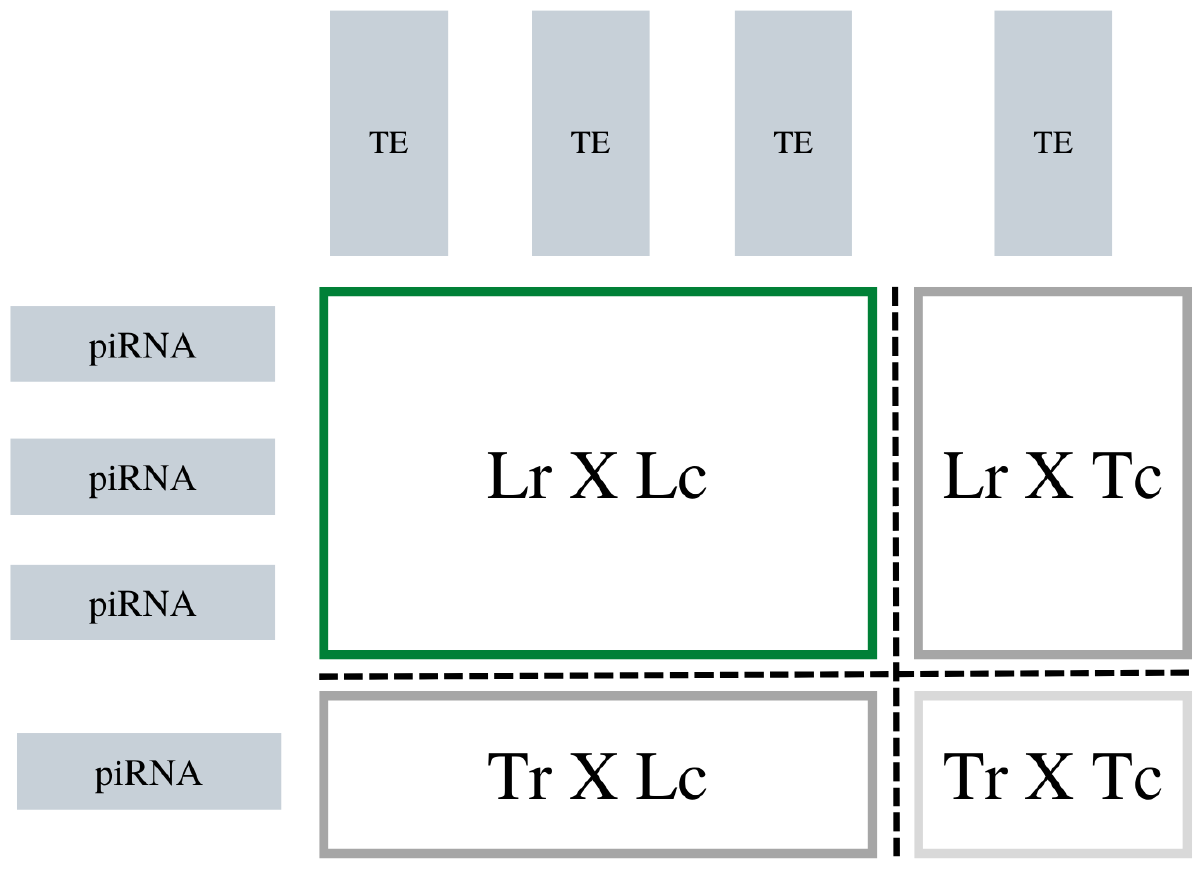
Interaction prediction scheme. Adapted from [36].

The global multiple-output approach (Figure 4(c)), inherent to the bi-clustering method proposed by [28], uses the sets of features *X*_*r*_ and *X*_*c*_ (row features and column features), so that the goal is to predict interactions between these sets. The process involves building a decision tree that incorporates these feature spaces so that the tree contains node tests for instances from both sets of interactions. The Y interactions matrix is vertically and horizontally partitioned. All features from both sets of instances are considered candidates for splitting at each new cut. The best cut is chosen by reducing the maximum impurity (variance) of Y. Given that *Φ*_*r*_ corresponds to the features of *X*_*r*_, that is, of piRNAs, and that *Φ*_*c*_ corresponds to the features of *X*_*c*_, that is, of TEs, the reduction of impurity is accumulated from row and column impurity reduction. If a test split in the tree is being applied on a piRNA feature, that is, in *Φ*_*r*_, the reduction is calculated by 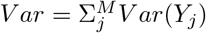, where M is the number of columns. If a test split in the tree is being applied on a TE feature, that is, in *Φ*_*c*_, the reduction is calculated by 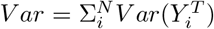, where N is the number of rows and *Y* ^*T*^ is the transpose matrix of Y. The process is illustrated in Figure 6.

**Fig 6.**
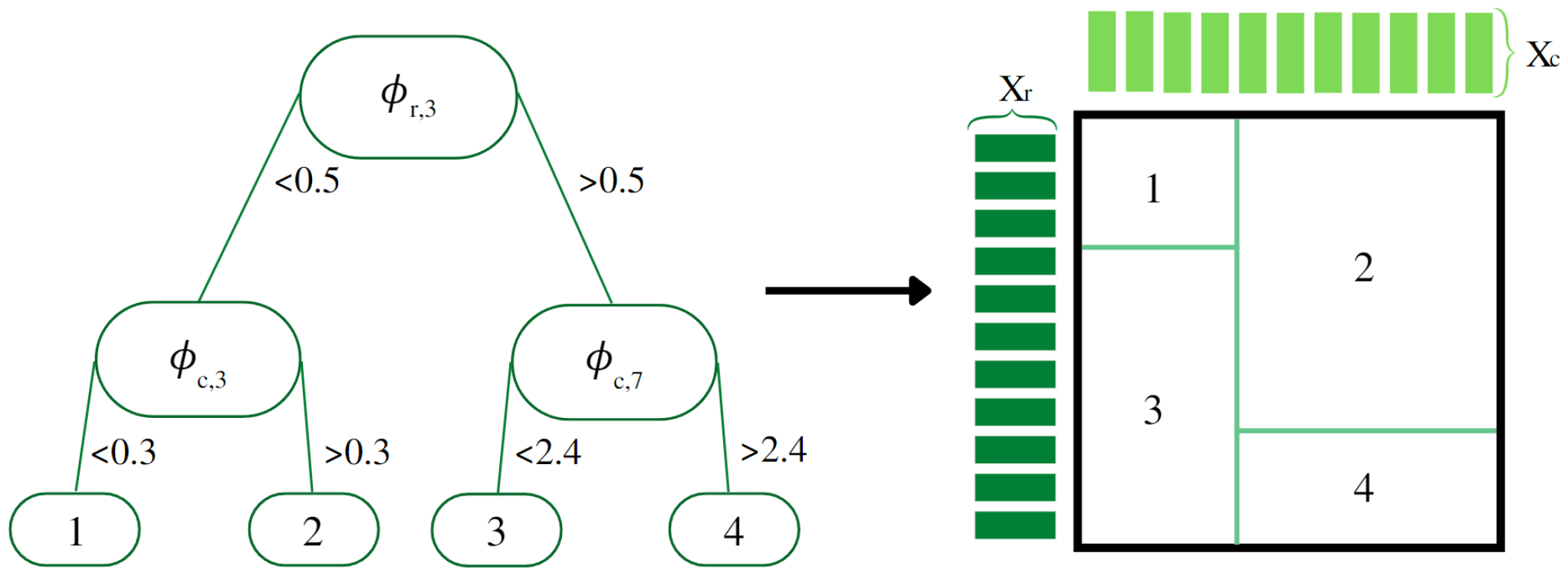
Illustration of how a bi-clustering tree works. On the left, the decision tree with *Φ*_*r*_ and *Φ*_*c*_, corresponding to the features of the row and column instances, respectively. On the right, the corresponding interaction matrix partitioned by the same tree. Adapted from [28, 33, 36].

### Cross-validation

According to [39], cross-validation is a statistical method for evaluating learning algorithms. The method separates the training and test data into successive rounds so that each data point can be validated (i.e., a data point must compose the training set at least once and compose the test set at least one time). The method employed in this study is k-fold cross-validation. In this method, data is divided into k segments of equal size so that, at each iteration, a different segment is used for testing, while the remaining k-1 is used for learning. In machine learning, cross-validation with ten iterations (i.e., with k = 10) is the usual.

Considering the approach of Pliakos *et al*. [28], we can see that each of the k iterations in a k-fold cross-validation experiment generates 1 training set and 3 test sets (Figure 5), resulting in a total experiment with k training sets and 3k test sets. In addition, the authors also recommend using k = 5 since, given the sparsity of data, 10-fold cross-validation can be a problem in the Tr × Tc setting, given the probability of having segments with zeros only.

### Evaluation metrics

When implementing a prediction model, the results are quantified by evaluation metrics that make the performances of any models comparable to each other. According to Baldi *et al*.. [40], the performances of classification prediction algorithms, as is the case of the present study, can be obtained in diverse ways. One of the canonical methods is the established by Rand [41], baptized Rand index, but simply called accuracy due to its ubiquitous use. The index, within the scope of decisions of an algorithm, can be given by 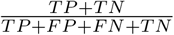, where TP are true positives, TN are true negatives, FP are false positives, and FN are false negatives, that is, the index tells the percentage of correct answers by the total.

However, in unbalanced learning, as in the case of the present study (we have 0.000377% of interaction pairs and 0.999623% of unknown pairs), the accuracy tends to be around 99% since the algorithms tend to classify according to the majority class, which is 0 in this data set. Thus, it is an uninformative method to assess the discernment capacity of the algorithm [42]. Thus, more informative alternatives were considered to evaluate the model.

According to Davis & Goadrich [43], the ROC (Receiver Operating Characteristic) curve is a common alternative, which shows how the number of correctly positive labeled examples varies with the number of incorrectly labeled examples as positive (that is, true positive rate over false positive rate). The true positive rate is given by 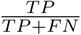,and the false positive rate is given by 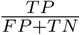. However, in highly unbalanced data sets, the ROC curve may present a more optimistic view of the algorithm since a significant change in the number of false positives has little effect on the false positive rate.

Given the highly unequal amounts of positive and negative interactions, using the PR curve (Precision-Recall) is the most indicated metric. The recall is given by 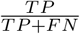(i.e., it is identical to the true positive rate), and the precision is given by 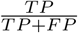. Precision, therefore, better represents changes in the number of false positives, as it compares false positives with true positives rather than true negatives [43].

While ROC and PR curves are good visual representations of the behavior of the algorithm, AUROC (Area Under the Receiver Operating Characteristic Curve) and AUPRC (Area Under the Precision-Recall Curve) are what effectively numerically inform model performance. The calculus of the area is carried out by numerical integration of the trapezoidal areas (trapezoidal rule) generated by each point of the ROC curve or the PR curve [43].

### Confusion matrix

According to [42], many machine learning algorithms return probabilities, not classes, as in the case of this study. In this sense, continuous non-binary values must be converted into binary (discrete) values based on decision thresholds so that all probabilities equal to or greater than the threshold are converted to 1, and those smaller than the threshold are converted to 0. For the ROC curve, the best threshold can be found using the G-mean (geometric mean). For this, the sensitivity and specificity of the algorithm are used. Sensitivity is equal to the true positive rate, conceptualized in the previous topic, and specificity is the complement of the false positive rate (1 - FPR), also conceptualized in the previous topic. The geometric mean is given by 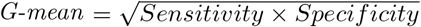. As for the PR curve, the best threshold can be found by the F-measure, given by 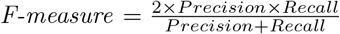.

The thresholds with the highest values for each measure are chosen as the best decision thresholds for the data set in question. For each threshold, one can build confusion matrices. A confusion matrix is merely a table that allows quantitatively verifying the algorithm’s measures of TP, FP, TN, and FN, comparing the predicted value with the actual value. However, since the number of tested thresholds is vast, only the confusion matrix for the best thresholds was calculated.

The confusion matrix was calculated by vectorizing the matrices, both the original interaction matrix and the predicted matrix (according to the best threshold). The vectorization guarantees that all the 13,841,400 items of the matrix are checked with no duplicates or missing values. The confusion matrices were computed solely to help understand the different behaviors of BICT and BICTR regarding positive interaction prediction (the primary interest in the study), as they are binary confusion matrices (target is 0 or 1). The standard evaluation metrics for a multi-label problem are AUROC and AUPRC, as already described.

## Results

### ROC curves

Table 1 shows the AUROC results obtained in the 5-fold cross-validation experiment, while Table 2 shows the AUROC results obtained in the 10-fold cross-validation experiment. The ROC curve of the BICTR method for one of the *Tr × Tc* folds is shown in Figure 7. The ROC curve of the BICT method for the same fold is shown in Figure 8.

**Table 1.** AUROC for the 5-fold cross-validation experiment.

| | $Tr \times Lc$ | | $Lr \times Tc$ | | $Tr \times Tc$ | |
| --- | --- | --- | --- | --- | --- | --- |
|  | BICTR | BICT | BICTR | BICT | BICTR | BICT |
| Fold 1 | 0.468 | 0.539 | 0.437 | 0.550 | 0.476 | 0.543 |
| Fold 2 | 0.474 | 0.532 | 0.448 | 0.535 | 0.478 | 0.511 |
| Fold 3 | 0.454 | 0.535 | 0.424 | 0.543 | 0.442 | 0.516 |
| Fold 4 | 0.456 | 0.533 | 0.474 | 0.531 | 0.479 | 0.517 |
| Fold 5 | 0.467 | 0.521 | 0.470 | 0.539 | 0.470 | 0.515 |
| Mean | 0.464 | 0.532 | 0.451 | 0.540 | 0.469 | 0.520 |

**Table 2.**
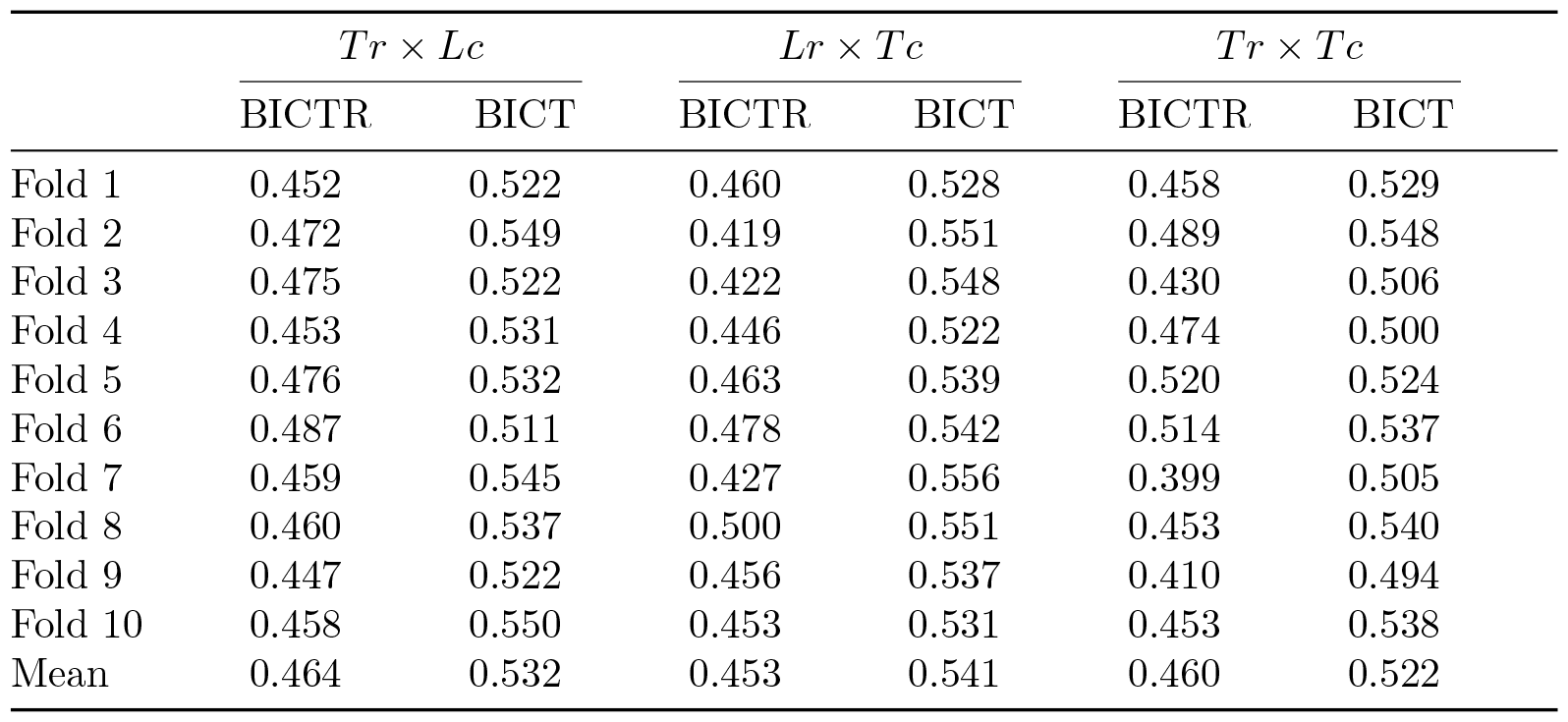
AUROC for the 10-fold cross-validation experiment.

| | $Tr \times Lc$ | | $Lr \times Tc$ | | $Tr \times Tc$ | |
| --- | --- | --- | --- | --- | --- | --- |
|  | BICTR | BICT | BICTR | BICT | BICTR | BICT |
| Fold 1 | 0.452 | 0.522 | 0.460 | 0.528 | 0.458 | 0.529 |
| Fold 2 | 0.472 | 0.549 | 0.419 | 0.551 | 0.489 | 0.548 |
| Fold 3 | 0.475 | 0.522 | 0.422 | 0.548 | 0.430 | 0.506 |
| Fold 4 | 0.453 | 0.531 | 0.446 | 0.522 | 0.474 | 0.500 |
| Fold 5 | 0.476 | 0.532 | 0.463 | 0.539 | 0.520 | 0.524 |
| Fold 6 | 0.487 | 0.511 | 0.478 | 0.542 | 0.514 | 0.537 |
| Fold 7 | 0.459 | 0.545 | 0.427 | 0.556 | 0.399 | 0.505 |
| Fold 8 | 0.460 | 0.537 | 0.500 | 0.551 | 0.453 | 0.540 |
| Fold 9 | 0.447 | 0.522 | 0.456 | 0.537 | 0.410 | 0.494 |
| Fold 10 | 0.458 | 0.550 | 0.453 | 0.531 | 0.453 | 0.538 |
| Mean | 0.464 | 0.532 | 0.453 | 0.541 | 0.460 | 0.522 |

**Fig 7.**
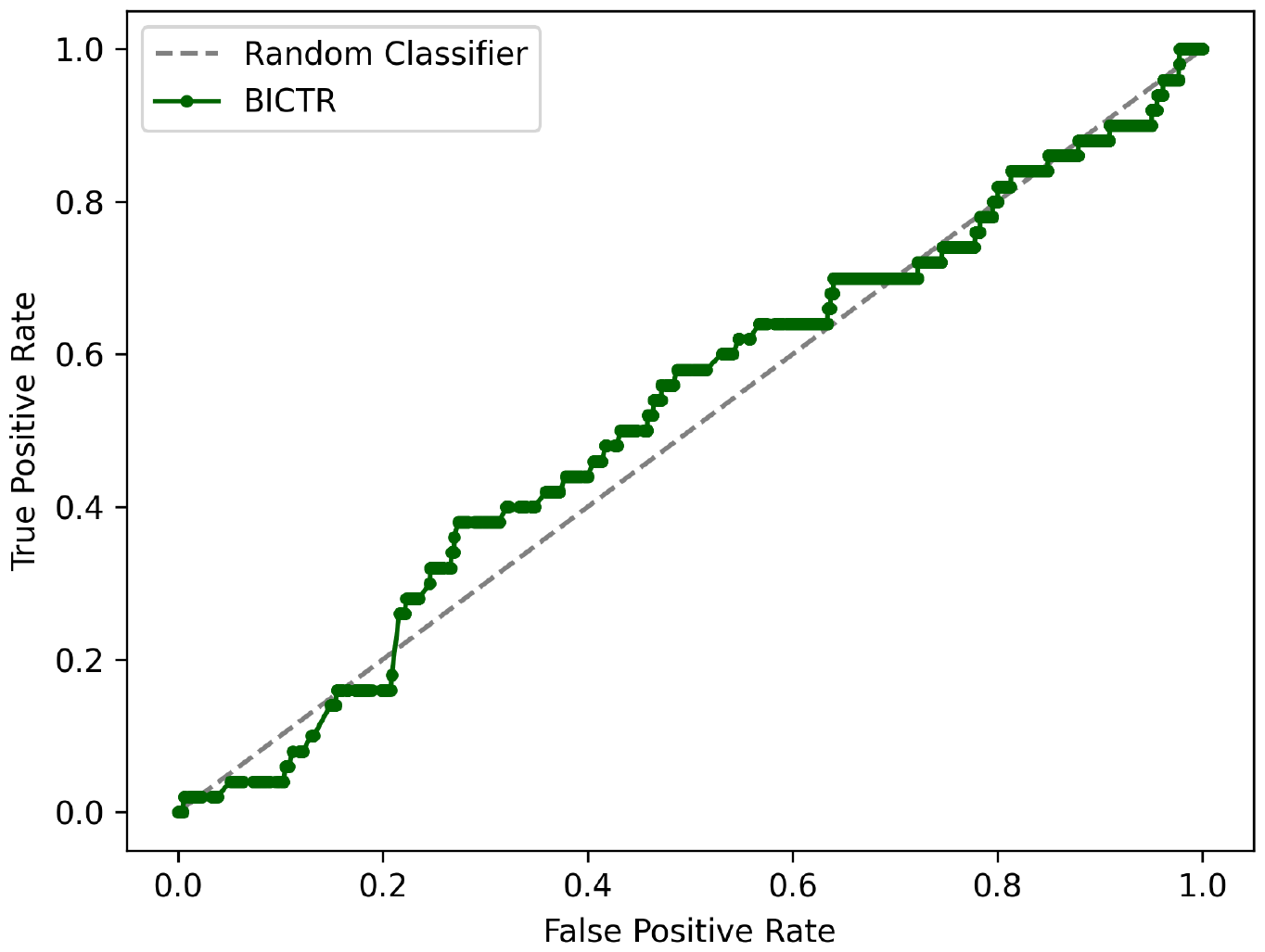
BICTR ROC curve for the *Tr × Tc* 5^th^ fold (10-fold).

**Fig 8.**
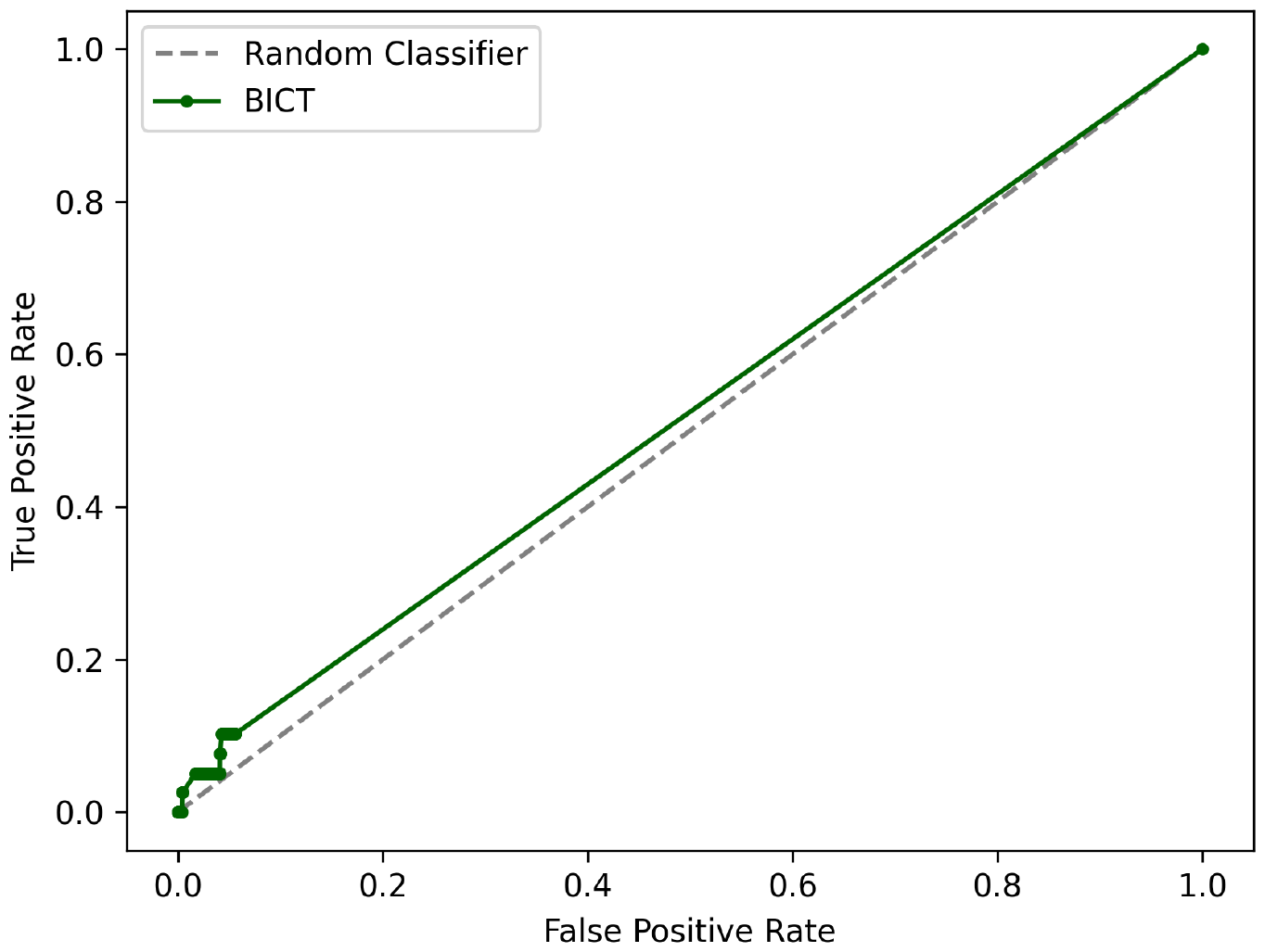
BICT ROC curve for the *Tr × Tc* 5^th^ fold (10-fold).

The confusion matrices for the best thresholds of that same fold, both for BICTR and the BICT, are shown in Figure 9. The best threshold was defined according to the highest G-mean value.

**Fig 9.**
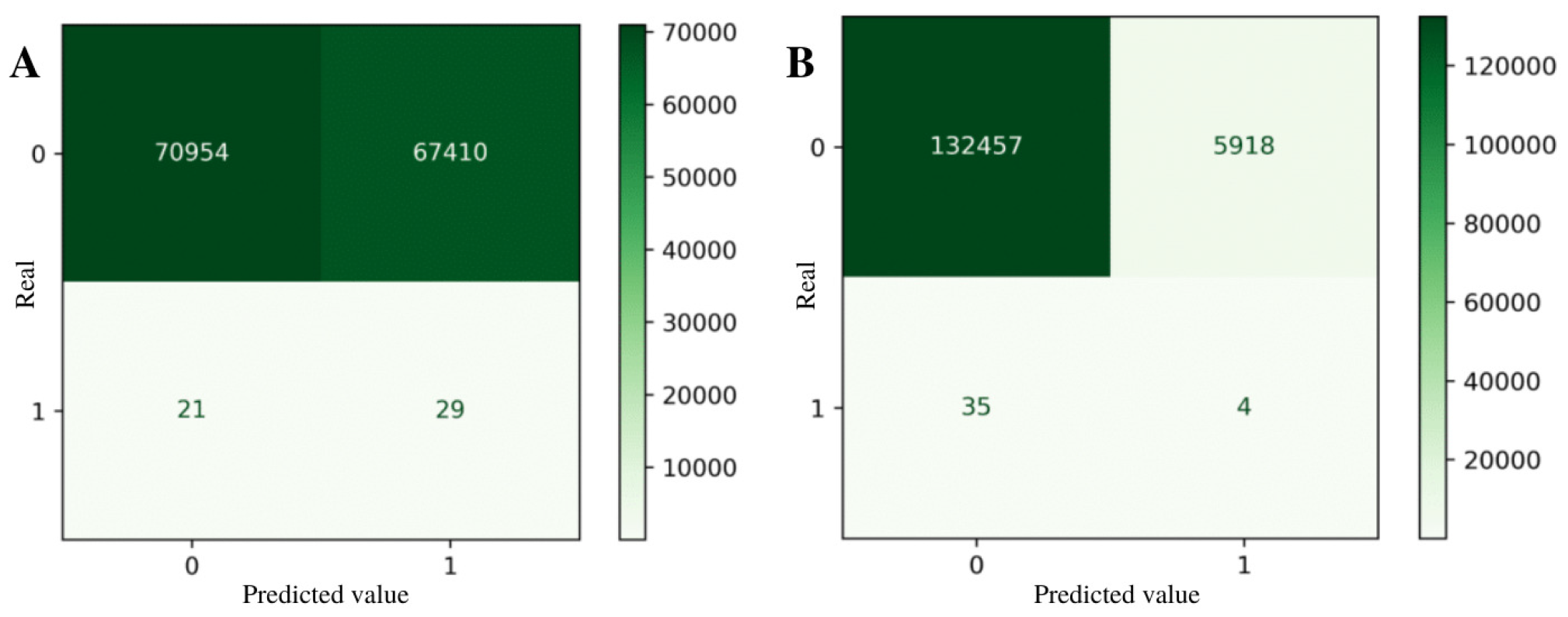
Confusion matrices for the best ROC curve thresholds for the *Tr × Tc* 5^th^ fold (10-fold): A) BICTR; B) BICT.

### PR curves

Table 3 displays the AUPRC results obtained in the 5-fold cross-validation experiment, while Table 4 shows the results of the AUPRC obtained in the 10-fold cross-validation experiment. The Precision-Recall (PR) curve of the BICTR for one of the *Tr × Tc* folds is presented in Figure 10. The PR curve of the BICT for the same fold is presented in Figure 11.

**Table 3.** AUPRC for the 5-fold cross-validation experiment.

| | $Tr \times Lc$ | | $Lr \times Tc$ | | $Tr \times Tc$ | |
| --- | --- | --- | --- | --- | --- | --- |
|  | BICTR | BICT | BICTR | BICT | BICTR | BICT |
| Fold 1 | $1.62 \times 10^{-3}$ | $7.03 \times 10^{-4}$ | $3.07 \times 10^{-4}$ | $1.03 \times 10^{-3}$ | $3.54 \times 10^{-4}$ | $1.02 \times 10^{-3}$ |
| Fold 2 | $1.61 \times 10^{-3}$ | $8.05 \times 10^{-3}$ | $9.15 \times 10^{-4}$ | $2.72 \times 10^{-3}$ | $2.82 \times 10^{-3}$ | $5.23 \times 10^{-4}$ |
| Fold 3 | $3.77 \times 10^{-4}$ | $2.37 \times 10^{-3}$ | $2.85 \times 10^{-4}$ | $9.40 \times 10^{-4}$ | $3.22 \times 10^{-4}$ | $5.22 \times 10^{-4}$ |
| Fold 4 | $3.84 \times 10^{-4}$ | $1.39 \times 10^{-3}$ | $1.54 \times 10^{-3}$ | $6.87 \times 10^{-4}$ | $3.47 \times 10^{-4}$ | $4.85 \times 10^{-4}$ |
| Fold 5 | $3.27 \times 10^{-4}$ | $5.82 \times 10^{-4}$ | $3.83 \times 10^{-4}$ | $3.47 \times 10^{-3}$ | $2.81 \times 10^{-3}$ | $6.01 \times 10^{-4}$ |
| Mean | $8.62 \times 10^{-4}$ | $2.62 \times 10^{-3}$ | $6.86 \times 10^{-4}$ | $1.77 \times 10^{-3}$ | $1.33 \times 10^{-3}$ | $6.29 \times 10^{-4}$ |

**Table 4.** AUPRC for the 10-fold cross-validation experiment.

| | $Tr \times Lc$ | | $Lr \times Tc$ | | $Tr \times Tc$ | |
| --- | --- | --- | --- | --- | --- | --- |
|  | BICTR | BICT | BICTR | BICT | BICTR | BICT |
| Fold 1 | $3.46 \times 10^{-4}$ | $1.70 \times 10^{-3}$ | $3.32 \times 10^{-4}$ | $7.45 \times 10^{-4}$ | $3.55 \times 10^{-4}$ | $1.46 \times 10^{-3}$ |
| Fold 2 | $3.32 \times 10^{-4}$ | $1.51 \times 10^{-3}$ | $1.52 \times 10^{-3}$ | $8.72 \times 10^{-4}$ | $3.67 \times 10^{-4}$ | $2.66 \times 10^{-3}$ |
| Fold 3 | $3.72 \times 10^{-4}$ | $2.90 \times 10^{-3}$ | $2.98 \times 10^{-4}$ | $1.15 \times 10^{-3}$ | $4.28 \times 10^{-4}$ | $3.55 \times 10^{-4}$ |
| Fold 4 | $3.41 \times 10^{-4}$ | $9.64 \times 10^{-4}$ | $1.29 \times 10^{-3}$ | $6.14 \times 10^{-4}$ | $4.63 \times 10^{-4}$ | $4.43 \times 10^{-4}$ |
| Fold 5 | $3.60 \times 10^{-4}$ | $2.87 \times 10^{-3}$ | $3.03 \times 10^{-4}$ | $7.51 \times 10^{-4}$ | $3.77 \times 10^{-4}$ | $4.36 \times 10^{-4}$ |
| Fold 6 | $3.85 \times 10^{-4}$ | $5.13 \times 10^{-4}$ | $3.55 \times 10^{-4}$ | $1.01 \times 10^{-3}$ | $3.58 \times 10^{-4}$ | $1.08 \times 10^{-3}$ |
| Fold 7 | $1.46 \times 10^{-3}$ | $1.03 \times 10^{-3}$ | $2.72 \times 10^{-4}$ | $2.53 \times 10^{-3}$ | $3.51 \times 10^{-4}$ | $4.55 \times 10^{-4}$ |
| Fold 8 | $3.41 \times 10^{-4}$ | $1.95 \times 10^{-3}$ | $1.96 \times 10^{-3}$ | $3.64 \times 10^{-3}$ | $6.28 \times 10^{-4}$ | $6.99 \times 10^{-4}$ |
| Fold 9 | $3.07 \times 10^{-4}$ | $5.46 \times 10^{-4}$ | $3.06 \times 10^{-4}$ | $2.65 \times 10^{-3}$ | $2.93 \times 10^{-4}$ | $4.48 \times 10^{-4}$ |
| Fold 10 | $3.54 \times 10^{-3}$ | $4.20 \times 10^{-3}$ | $1.31 \times 10^{-3}$ | $7.33 \times 10^{-4}$ | $4.19 \times 10^{-4}$ | $9.61 \times 10^{-4}$ |
| Mean | $7.78 \times 10^{-4}$ | $1.82 \times 10^{-3}$ | $7.95 \times 10^{-4}$ | $1.47 \times 10^{-3}$ | $4.04 \times 10^{-4}$ | $9.00 \times 10^{-4}$ |

**Fig 10.**
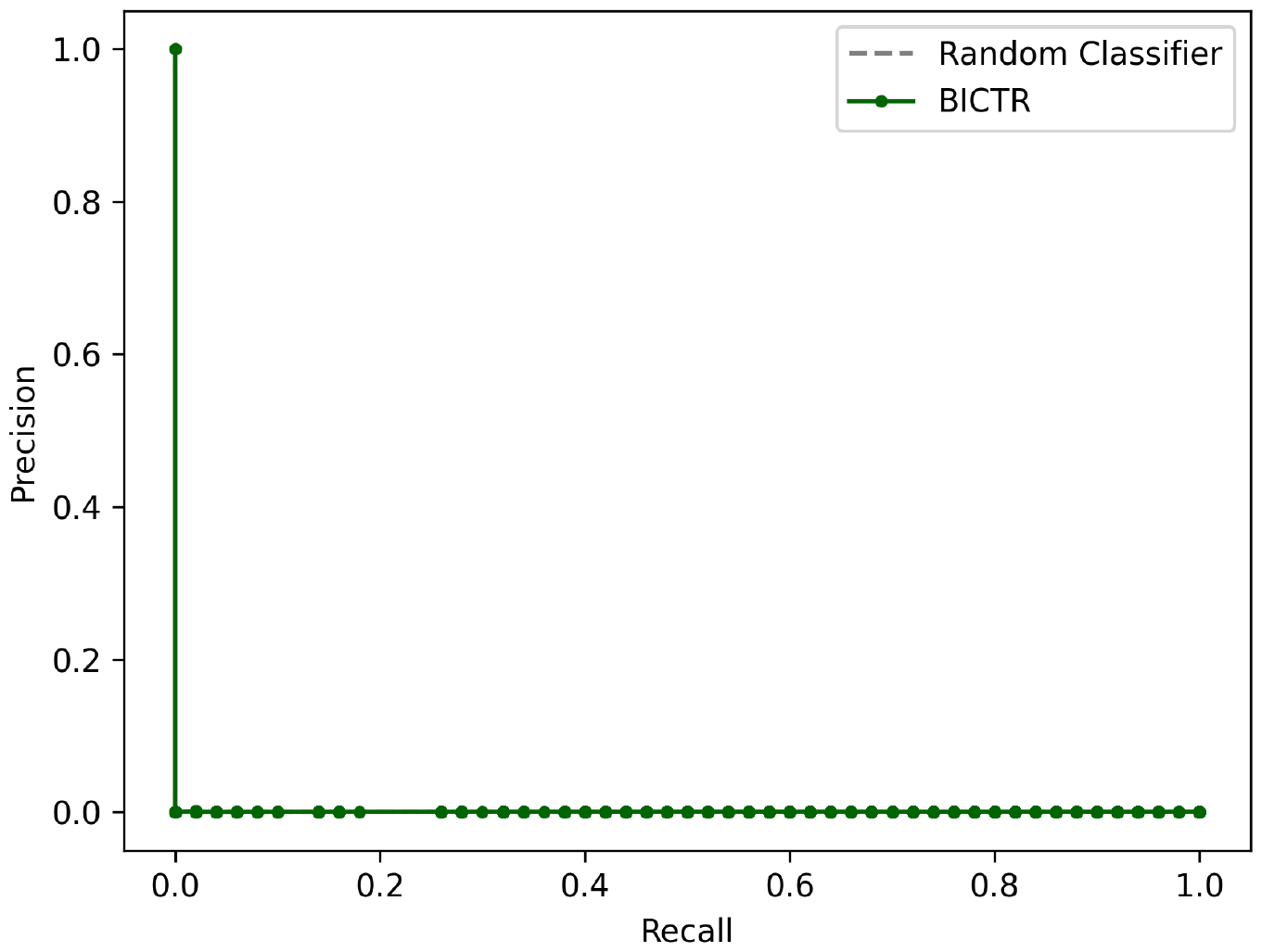
BICTR Precision-Recall curve for the *Tr × Tc* 5^th^ fold (10-fold).

**Fig 11.**
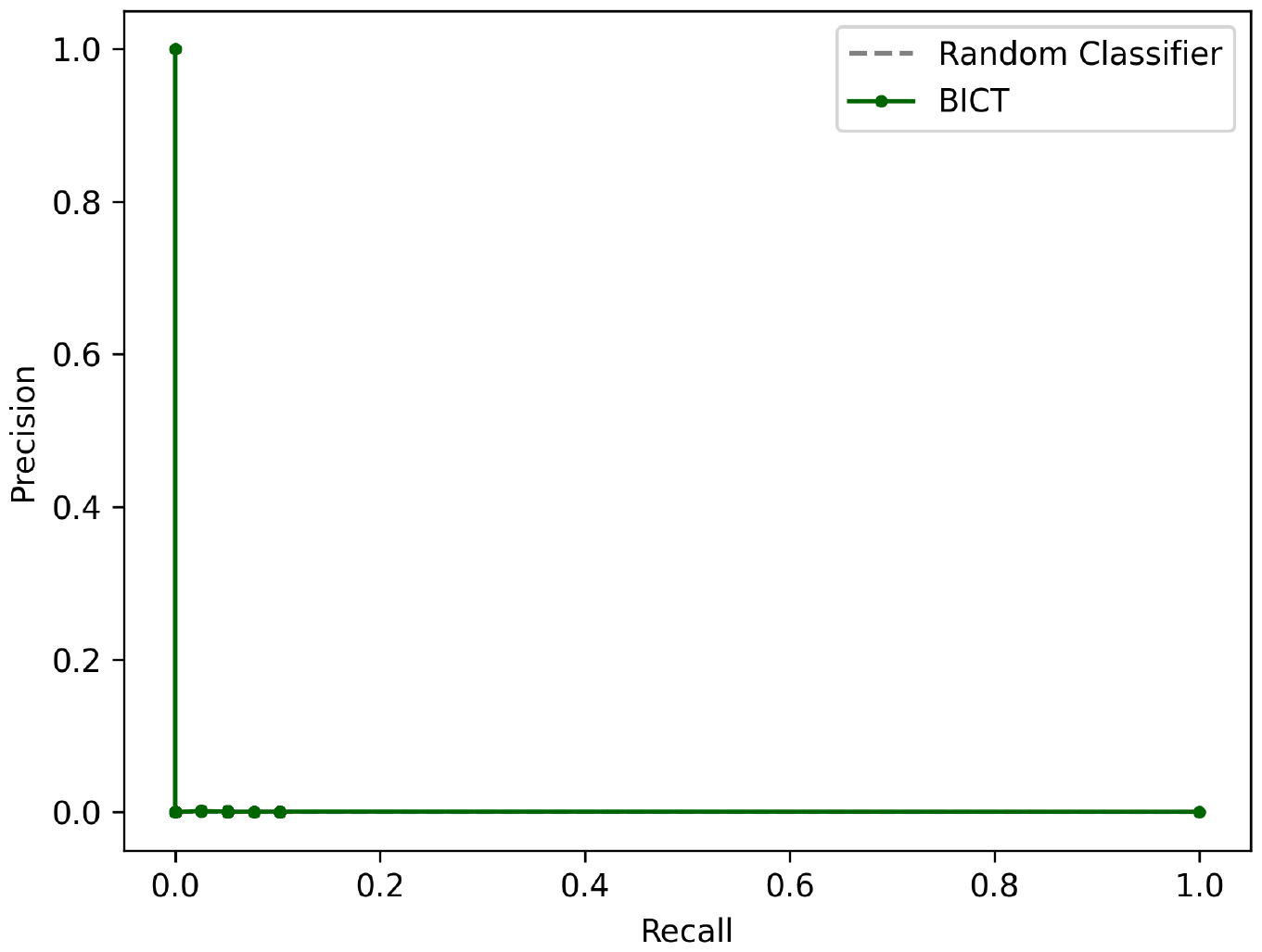
BICT Precision-Recall curve for the *Tr × Tc* 5^th^ fold (10-fold).

The confusion matrices for the best thresholds of that same fold, for both BICTR and the BICT, are represented in Figure 12. The best threshold was defined according to the highest F-score value.

**Fig 12.**
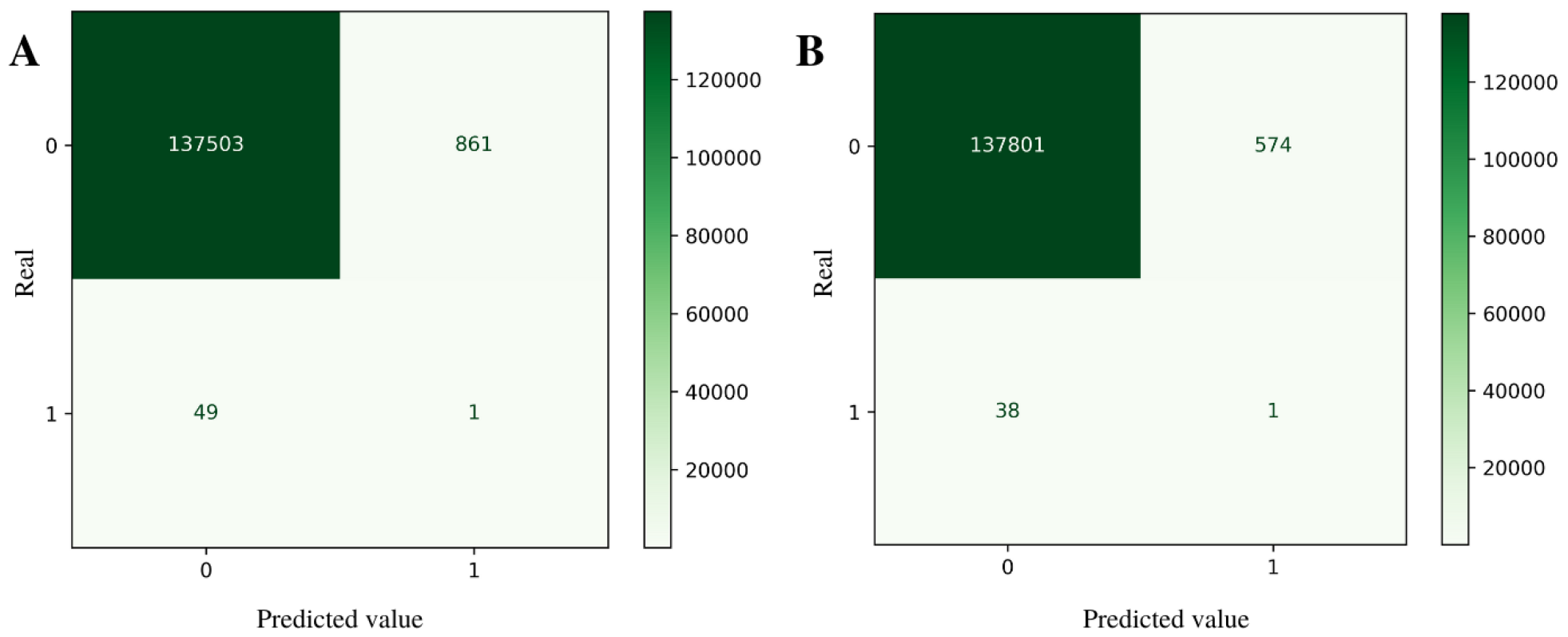
Confusion matrices for the best PR curve thresholds for the *Tr × Tc* 5^th^ fold (10-fold): A) BICTR; B) BICT.

## Discussion

### Learning results

Bearing in mind that, both for AUROC and AUPRC, the better the model, the closer the areas are to 1, it is evident from the values displayed in the tables that the algorithm was not able to predict the true interactions satisfactorily. However, it did correctly predict some positive ones, as evidenced in the confusion matrices of Figures 9 and 12. Observing the ROC charts, one can see that the behavior of the curve traced by the points is very similar to the behavior of a random classifier, oscillating around the line depicting it. By observing the PR charts, it is noticeable that the traced curve also resembles the curve of a random classifier. By means of these observations, it can be argued that the model for the present data did not obtain good learning and behaved as a random classifier, not being able to predict new interactions in such a satisfactory fashion. The massive amount of unknown/negative interactions (approximately 0.999623%) makes the prediction problem an extremely unbalanced classification problem, which makes learning the model challenging.

Comparing BICT and BICTR methods, BICTR, although used to improve the prediction performance, obtained smaller values of AUPRC and AUROC in most tasks. What happened can be understood by the confusion matrices. The factorization, indeed, increased the probability of classifying an interaction pair as 1. As a matter of fact, the values are very close to a 1:1 prediction ratio. However, although it slightly increased the TP score, it vastly increased the FP score (more than 11*×*), as can be seen in the confusion matrices of Figure 9.

In terms of effects in the laboratory, considering that all positively labeled interaction pairs would be experimentally tested, if using the BICTR output as a map of interactions to experiment, there would be a gain of more than 7 *×* of real positive interactions, compared to what would be yielded if the positively labeled output of BICT was tested. Even though it would still be necessary to test many pairs, the method applied drastically reduces the investigation space compared to if all the molecule interactions were to be tested. Therefore, although BICTR obtained smaller areas under the curves, the factorization may still ease the lab’s work, given that positive interactions are the actual interest in this problem.

One can also see that the BICTR, both in ROC and PR curves, obtained a smoother curve than the BICT curve. That happens because the curve is generated based on the interpolation of conversion thresholds, which are all the values in the predicted matrix (ranging from 0 to 1). Since, in BICT methodology, PBCT receives the original data, which contain many 0 values, many 0 values are predicted, limiting the number of threshold values used to plot the curve. Meanwhile, in BICTR, PBCT learns with repopulated data so that 0 values are not predicted, making the number of thresholds exceptionally larger and, consequently, the number of points plotted in the charts. With that in mind, it is worth noting that, although BICT presents larger areas than BICTR areas in the vast majority of cases (Tables 1 to 4), the smaller amount of BICT points interpolated (due to less conversion threshold values) causes greater gaps between the points, which contributes to a larger area but is not as trustworthy as the curve traced by BICTR, which is generated with many more points.

## Alternative approaches

According to [33] and Figure 4, besides bi-clustering trees, there are other approaches to deal with network inference problems: the global single output approach (GLSO) and the local multiple output approach (LOCMO).

To settle the extremely unbalanced data set issue, one can use sampling, both by oversampling and undersampling, since a good part of the classification algorithms are built in a way to operate better in data with equal numbers of observations for each class [42]. A classic oversampling technique example is SMOTE (Synthetic Minority Oversampling Technique), which uses synthetic data from a defined neighborhood, data which are created by interpolating several instances of the minority class so that the algorithm, instead of using the original data points, uses data attributes and their relationships [44].

In order to apply the global single output approach, the piRNA and TE feature vectors were concatenated to form only rows (unlike PBCT, where rows were dedicated to piRNA features alone), and a single binary column represented the presence or absence of an interaction between the corresponding piRNA-TE pair. In addition, SMOTE was applied. After this technique, there were equally 50% **0** and **1** data points. With post-SMOTE data, decision trees and random forest algorithms were applied. The random forest algorithm presented better results, according to Table 5.

**Table 5.** Global single output approach.

| Metric | DT | RF |
| --- | --- | --- |
| Precision | 0.0007 | 0.3896 |
| Recall | 0.2988 | 0.0347 |
| F-measure | 0.0014 | 0.0637 |

In order to apply the local multiple output approach, two distinct matrices were used, one with the piRNA features and the interaction matrix aside and another with the TE features and transposed interactions matrix aside. With these two matrices, a random forest classifier chain algorithm was applied. The results are presented in Table 6.

**Table 6.** Local multiple output approach.

| Metric | piRNA | TE |
| --- | --- | --- |
| Precision | 0.4998 | 0.8998 |
| Recall | 0.4999 | 0.5016 |
| F-Measure | 0.4990 | 0.5031 |

LOCMO presented better results, as can be seen in the F-measures. Dealing with very particular matrices once at a time resulted in higher precision and recall values, reflected in higher F-measures. The types of molecules, i.e., piRNAs and TEs, are physicochemically very different, and as the algorithms use those physicochemical features to predict the interaction, dealing with only a kind of molecule per prediction task could be beneficial. Molecule-molecule interaction prediction tasks usually rely on pairs of molecules with great affinity, which is not the case in piRNA-TE interaction, as exposed.

## Conclusion

To conclude, the relevance of this paper could be summed up as the potential to drastically reduce labor in molecular biology laboratories in the quest for finding molecule-molecule interactions. Even though PBCT was not satisfactory in our proposed method, many things can be tweaked in order to seek better results, such as different feature generation methods or the continued update of the model with more *in vivo* verified interactions, as the known interactions are still very few compared to unknown, as evidenced. While many trials could still be needed to achieve excellence in piRNA-TE prediction tasks, which could be addressed in future work, this pursuit was strongly justified in this paper.

## Acknowledgments

We thank Mr. Shen, Mr. Chen and Mr. Mello for providing the CLASH interaction data used in this paper.

